# The role of EPS in the selective biosorption and desorption of REEs

**DOI:** 10.64898/2026.07.02.736058

**Authors:** Memphis J. Hill, Brandon R. Briggs

## Abstract

Rare earth elements (REEs) are critical components of green technologies, but current mining and purification methods remain environmentally unsustainable due to their high energy consumption and intensive chemical requirements. Bio-hydrometallurgical processes have the potential to concentrate and recover REEs at a circumneutral pH. Work presented here uses bacteria at neutral pH to concentrate REEs from solution and subsequently recover those REEs using sodium citrate. *Shewanella oneidensis* MR-1 was incubated anaerobically in a culture media solution spiked with 14 REEs and yttrium for one to six days. REE concentrations remaining in solution were then compared to REE concentrations on cell pellets. For these same timepoints, the loosely bound extracellular polymeric substance (LB-EPS) was removed from cells prior to quantifying REEs on pellets to narrow down the location of REE binding. Moreover, cell pellets collected after 5 days in REE spiked solution were subjected to a time series desorption assay using sodium citrate. *Shewanella oneidensis* at a starting OD600 of 0.6 adsorbed 1.18mg/g of REE after 3 days. 80% of these REEs were located in the LB-EPS. In 10 minutes, 0.5 M sodium citrate desorbed about 75% of REEs from cells and over 95% after 24 hours. This method was also applied to Alaskan coal and showed that 68-86% of REEs were desorbed form S. oneidensis. This study elucidates the REE binding location and capacity of *S. oneidensis*, REE removal efficiency of sodium citrate overtime, and the application of this sustainable biotechnology for REE recovery at a circumneutral pH from Alaskan coal.

**Importance:** Rare earth elements (REEs) are essential for technologies such as electric vehicles, wind turbines, batteries, and medical devices, but current methods for obtaining them are energy-intensive and can harm the environment with acidic waste. This study demonstrates a non-acidic sustainable approach using naturally occurring bacteria to capture REEs and then recover them with a biologically derived solution. The research also identifies where these elements attach to the bacteria, providing new insight into how the process works and how it can be improved. Importantly, the method successfully recovered REEs from Alaskan coal, highlighting its potential for real-world applications. By improving our understanding of microbial methods for rare earth element recovery, this work could help reduce the environmental impacts of mining and support more sustainable, secure supplies of the critical materials needed for clean energy and other advanced technologies.

## 1. Introduction

Rare earth elements (REEs) encompass a group of 17 chemical elements, including the lanthanides, scandium and yttrium. REEs are indispensable in a wide breadth of economic sectors, including renewable energy, automobiles, medical science, electronics, manufacturing, and the military as key components in many current and emerging technologies.^1–4^ However, the extraction and recovery of REEs remain challenging because they are typically dispersed in low-grade ores and dilute solutions, and conventional recovery processes are both economically costly and environmentally damaging. As a result, biological approaches are being actively explored as sustainable alternatives for REE extraction and processing.^5–10^

*Shewanella oneidensis* MR-1 is one potential candidate for biomining technologies as it is known to extract and biosorb a wide array of metals including REEs.^11–17^ Recent studies have demonstrated the practical potential of this organism; for instance, Sachan et al. (2023) showed that *S. oneidensis* MR-1 can extract REEs from Alaskan coal at circumneutral pH, in some cases outperforming traditional sulfuric acid leaching with individual recovery rates as high as 96% for lanthanum (La).^14^ Furthermore, the genetic mechanisms for these interactions are beginning to emerge. Mendin et al. (2023) utilized a genome-wide knockout screen of *S. oneidensis* MR-1 to identify 242 genes controlling REE biosorption, revealing that cell-surface polysaccharides, pilus assembly, and redox regulatory systems strongly influence REE binding and selectivity, offering engineering targets for improved microbial separation of individual lanthanides.^15^ Specifically, they showed that operons involved in making O-antigens on the lipopolysaccharide layer increased biosorption of europium (Eu), while disruptions in the mannose-sensitive heme agglutination (MSHA) operon, a gene which synthesize pili on the outer membrane of *S. oneidensis* MR-1, reduced Eu-biosorption.

The genetic determinants identified above are typically associated with extracellular polymeric substances (EPS), which provide an array of metal-binding ligands. Functional groups on the cell wall and EPS, such as carboxyl, phosphate, and hydroxyl groups, form stable complexes with metal ions.^18,19^ These polar functional groups can bind metals, leading to the accumulation of metal ions on the bacterial surface, and potentially concentrating REEs from dilute solutions. Although *S. oneidensis* was previously believed to lack an EPS,^20,21^ FTIR spectra have since shown two distinct layers of EPS, loosely bound EPS (LB-EPS) and tightly bound EPS (TB-EPS).^12^ FTIR spectra from Yan et al. (2021) revealed stronger stretching vibrations from carboxyl and phosphate groups in the LB-EPS than the TB-EPS and found 4.7 times as many carboxyl groups and 2.4 times as many phosphate groups in the LB-EPS than in the TB-EPS of *S. oneidensis*.^12^ Although these distributional differences suggest a high degree of surface tunability, there is a lack of comprehensive understanding of *S. oneidensis* binding capacity, element selectivity, and physical location of adsorbed REEs.

In addition to unknowns for REE absorption, the efficient recovery of these elements from this biological matrix remains a critical knowledge gap for effective biorecovery of REEs. To address this recovery challenge, sodium citrate presents a promising candidate for metal desorption due to is chelating properties and as a biodegradable alternative to traditional metallurgical stripping agents.^22–24^ For example, Yan and Chen (2024) conducted four sequential desorptions of REE from *Escherichia coli* using 0.05 mM citrate at a pH of 6. The majority of REEs were desorbed in the first round, 50 to 200 ug/g of Y and REEs (Ce through to Er) and 300-500 ug/g of Yb and Lu. Interestingly, the maximum amount of Y (∼700ug/g) was desorbed in the third citrate wash.^25^ Citrate is composed of three carboxyl groups that can form stable, coordinate-covalent complexes with trivalent REEs.^26,27^ Unlike mineral acids which can denature the biomass and complicate downstream processing, sodium citrate is a mild, biodegradable, and biocompatible reagent.

This study investigates the REE binding capacity of *S. oneidensis* MR-1, the location of REE binding, and the potential of sodium citrate as a desorbing agent. We examine how incubation time influences both biosorption and desorption and compared recovery between heavy and light REEs. Finally, we apply this biotechnological approach to an REE-containing Alaskan coal feedstock. These analyses provide insights into the mechanisms governing REE biosorption and desorption and inform the development of sustainable bio-based strategies for REE recovery, separation, and environmental remediation.

## 2. Materials and Methods

### 2.1 Cell Culture

A 1 ml freezer stock vial of *S. oneidensis* MR-1 was thawed and used to inoculate 50 ml of Luria-Bertani (LB) medium containing yeast extract (5.0 g/L), sodium chloride (10.0 g/L) and tryptone (10.0 g/L). The inoculated broth was kept at 28 °C and shaking at 150 rpm for 24 hours. After 24 hours, the total culture volume was increased to 220 mL and left shaking for an additional 24 hours.

### 2.2 REE Binding

*S. oneidensis* in LB was collected at an OD600 of 0.945 +/- 0.03 in 50 mL falcon tubes and centrifuged for 10 minutes at 6000 rpm (5911xg) in a Sorvall Evolution RC centrifuge with SLA-600TC rotor. Cells were washed twice with Shewanella Minimal Media (SHMM) containing 29.76 mM sodium bicarbonate, 10mM PIPES, 1.34 mM KCl, and 20mM sodium-L-lactate. After washing the cells were resuspended in SHMM with 50ppb CMS-1 (a standard solution of REEs from Inorganic Ventures containing equal amounts of Ce, Dy, Er, Eu, Gd, Ho, La, Lu, Nd, Pr, Sc, Sm, Tb, Tm, U, Y, and Yb) for a total volume of 220 mL and starting OD600 between 0.6 and 0.7. All media was autoclaved at 121°C for 20 minutes prior to use. Na-L-lactate was sterilized through a 0.2µm filter and 20mM was added to the autoclaved medium. CMS-1 was also added after autoclaving for a final concentration of 50 ppb of each element. Promethium was not tested as it does not occur naturally in the Earth’s crust.^28^

Aliquots of 50ml of cell culture were set up for each timepoint (days 1 through 6) and left shaking in acid-washed glass bottles in a 30°C incubator. For each time point, we collected triplicate technical replicates to measure OD600, dry pellet weights, REEs bound, REE remaining in culture media, and REEs remaining on pellet after LB-EPS removal.

### 2.3 Extracellular Polymeric Substance Removal

The LB-EPS was separated from cell pellets following a heat treatment protocol described by Yan et al. (2021). Triplicate samples were collected from each culture condition and spun at 6,000 x g for 10 minutes at room temperature. Supernatants were removed and saved for REE quantification via ICP-MS. Each pellet was resuspended in 4 ml of room temperature 0.9% NaCl and then further diluted with 3 ml of 0.9% NaCl at 70°C. Resuspended pellets were then capped tightly and vortexed horizontally on a Vortex-Genie for 2 minutes. Cells were then centrifuged at 5000 x g for 15 minutes at room temperature. The supernatants were decanted into new tubes and filtered through a 0.22 µm syringe filter. Filtrates containing the LB-EPS were stored at -20°C. The pellets remaining in the centrifuge tubes were acidified and diluted with 2% HNO_3_ for REE quantification via ICP-MS.

### 2.4 Optical Density Measurements

For OD600 measurements, 600 µl of cells were collected from each 50ml cell culture condition. That 600 µl was then split to 200 µl per well in a 96-well flat bottom plate for triplicate OD600 measurements using a BioTek Cytation 1 cell imaging multimode plate reader.

### 2.5 Dry Pellet Weights

Triplicate cell samples from 3 ml of culture were pelleted via centrifugation for 10 minutes at 6000 rpm (5911xg) in a Sorvall Evolution RC centrifuge with SLA-600TC rotor. Supernatants were removed and saved for measurements of REEs remaining in the culture media. Empty 1.5 ml Eppendorf centrifuge tubes were pre-weighed for each replicate for each time point. Each cell pellet was resuspended in 1ml of nanopure MilliQ water for a ratio of 3 ml of cell culture per 1 ml of water, transferred to a 1.5 ml Eppendorf centrifuge tubes and spun at 10,000 x g for 10 minutes in an Eppendorf 4518 centrifuge with FA-45-18-11 rotor. The water rinse was saved for ICP-MS and then pellets in the pre-weighed tubes were resuspended in 2% HNO_3_ for triplicate measurements of total REEs bound to cells. After ICP-MS, the acidified pellets were spun again at 10,000 x g for 10 minutes, and the pre-weighed tubes were left uncapped in a 30°C incubator overnight to dry and then weighed again. The difference in weight between the empty tube and the tube containing the dried pellet was used to determine dry pellet weight.

### 2.6 REE Desorption Assay

For the desorption time course, cells were grown as described in section 2.1 and incubated in SHMM with 50 ppb REEs in acid washed glass bottles capped to limit oxygen and kept at 30 °C shaking at 150 RPM for just under 6 days (5 days and 18 hours). Cells were pelleted in 50 mL falcon tubes for 10 minutes at 6000 rpm in a Sorvall Evolution RC centrifuge with SLA-600TC rotor and washed once with nanopure MilliQ water. Cell pellets were resuspended in either 0.5M sodium citrate dihydrate (Na-citrate) from J.T Baker® or nanopure MilliQ water. Cells were concentrated into 1 ml of resuspension liquid per 3 ml of original cell culture. 1 ml aliquots were made in triplicate for Na-citrate desorption times of: 10, 20, 30, 40, 50, 60, and 90 minutes; 2, 3, 4, 5, 6, 12, 24, and 48 hours. Cells resuspended in MilliQ nanopure water for triplicate negative controls were incubated for 0.5, 3, 24, and 48 hour time points. To ensure no contamination with cells, the sodium citrate supernatants were centrifuged again prior to aliquoting and acidifying for ICP-MS. REEs in sodium citrate solution and REEs remaining on cells after each incubation period were quantified via ICP-MS following steps described in section 2.7.

### 2.7 ICP-MS

All REE concentrations were determined by ICP-MS from aqueous solutions of 2% nitric acid. Supernatants were acidified to 2% HNO_3_ with 70% trace metal-grade nitric acid and then diluted 1:150 in 2% HNO_3_. Cells were pelleted for 5 minutes at 10,000 x g in an Eppendorf 4518 centrifuge with FA-45-18-11 rotor and resuspended in 1ml of 2% HNO_3_. The cells were then briefly vortexed before being diluted 1:40 in 2% HNO_3_ for analysis on an Agilent 7500c ICP-MS. Due to the very low amounts of REEs either remaining on cells after Na-citrate desorption or desorbed by the water-only control conditions, several samples were rerun on the ICP-MS at only a 1:10 dilution in 2% HNO_3_. A calibration curve was constructed using CMS-1 at known concentrations of 100, 50, 10, 1, 0.5, 0.1, and 0.01 ppb. Quality controls of CMS-1 at 10 ppb and 0.5 ppb were run every 10 samples to test for instrument drift, and all elements except Sc, U, and Th were consistently detected throughout the run. An internal standard of 1000 ppb Rhenium and Indium in 0.1% Triton-X was used throughout the run. All results reported had concentration RSDs less than 15.

### 2.8 Coal Incubations

Coal was crushed and sieved to a diameter of 1.19 mm or less and then autoclaved to sterilize. Glass serum bottles were acid washed and autoclaved. 1.4 grams of sterile coal (10% w/v) were weighed into triplicate serum bottles and then inoculated with SHMM containing 20mM lactate and *S. oneidensis* MR-1 at an OD600 of 0.8. Abiotic controls received the same amount of coal and culture media, but no bacteria. Serum bottles were sealed with a rubber stopper and aluminum clamp then set to shake at 120 rpm for 7 days. On the fourth day, cultures were off gassed and then 60 ml of air was sterilely injected through a 0.2 µm filter into each bottle. On the seventh day, the seals were broken, and the coal-media-cells mixtures were removed from the serum bottles. These mixtures were then centrifuged at 6000 rpm for 10 min and supernatant was collected in a clean 50ml conical. The cells were then separated from the coal as described below.

### 2.9 Nycodenz separation of cells from coal

Coal-cell mixtures were resuspended in 3ml of nanopure water and then diluted 1:2 with wash buffer that per ml contained 600 µl of 2.5% NaCl, 200 µl of 1%(v/v) Tween 80, and 200 µl of methanol. The mixture was briefly vortexed and then sonicated in a water bath for three minutes. Nycodenz cushions were prepared in clean 15 ml conical tubes with 1.5 ml of 45% Nycodenz solution. The resuspended coal-cell mixtures were then gently dispensed on top of the Nycodenz cushion, being careful not to mix. The coal-cell resuspensions layered on top of Nycodenz cushions were then centrifuged for 20 minutes at 4000 rpm in an Eppendorf Centrifuge 5810 R with a swinging bucket rotor without the brake. A sterile transfer pipet was then used to move the cells to a fresh 15 ml tube where they were diluted 1:3 with nanopure water and then centrifuged at 6,000 x g for 10 minutes to pellet the cells. The supernatant was decanted to remove any remaining Nycodenz. Cells and supernatants were processed for ICP-MS as described in section 2.7.

### 2.10 Equations

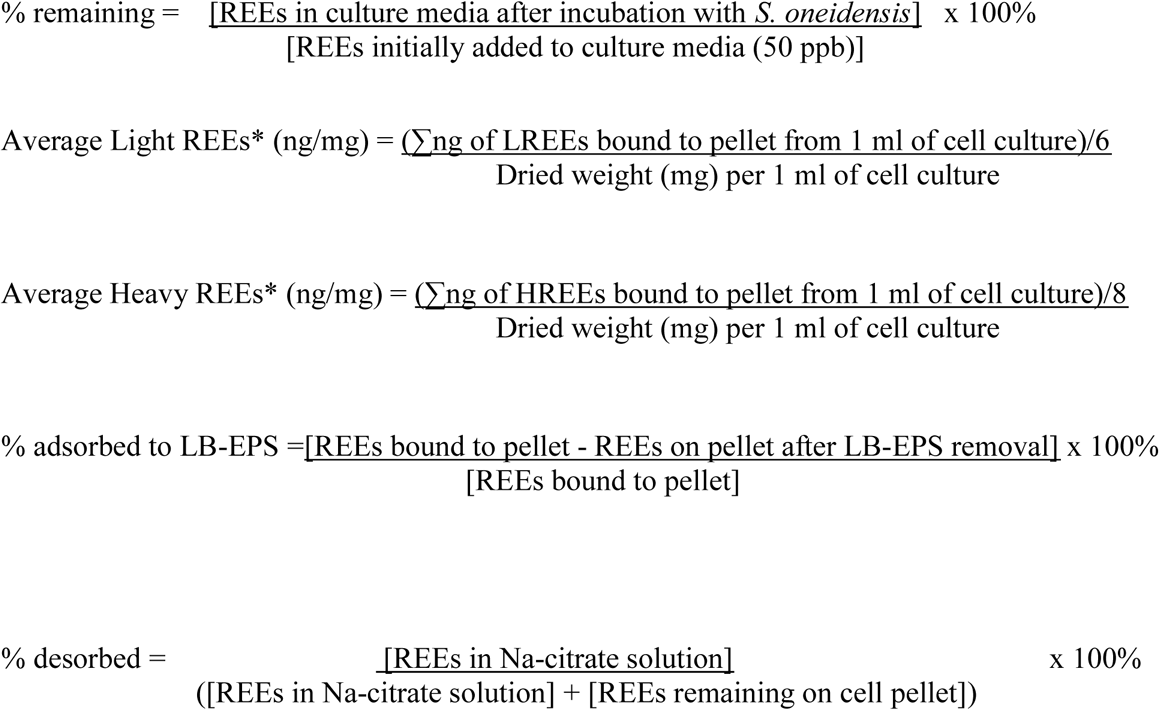

*Light REEs are La, Ce, Pr, Nd, Sm, and Eu. Heavy REEs are Y, Gd, Tb, Dy, Ho, Er, Tm, Yb, and Lu.

Loss of some REEs in the abiotic controls over time (likely due to adsorption to the glass bottles) was noted. The percent of REEs remaining in media and lost during processing or to adsorption to the container in abiotic conditions on day 0 and 5 are shown in the Supplementary Information (Figure SI-1).

## 3. Results and Discussion

### 3.1 REE Biosorption

The REE adsorption capacity of *S. oneidensis* MR-1 in anoxic conditions was tested for 14 lanthanide elements and yttrium (Y). The average REE adsorbed per dry weight of cells increased until Day 3 for both light and heavy REEs (Figure 1). On Day 3, light REEs were adsorbed to cells significantly (p = 0.009) more than heavy REEs with 88% and 73% adsorption, respectively. This equates to 500 ng of light REEs and 650 ng of heavy REEs adsorbed per mg of cell dry weight. Adsorption decreased after Day 3, with Day 5 also having significantly different adsorption between light and heavy REEs (p = 0.006). For both heavy and light REEs, there was significantly more adsorption on Day 3 compared to Day 1 (p = 0.007 for lights and p = 0.008 for heavies) and significantly less adsorption on Day 6 than on Day 3 (p = 0.008 for lights and p = 0.002 for heavies).

**Figure 1.**
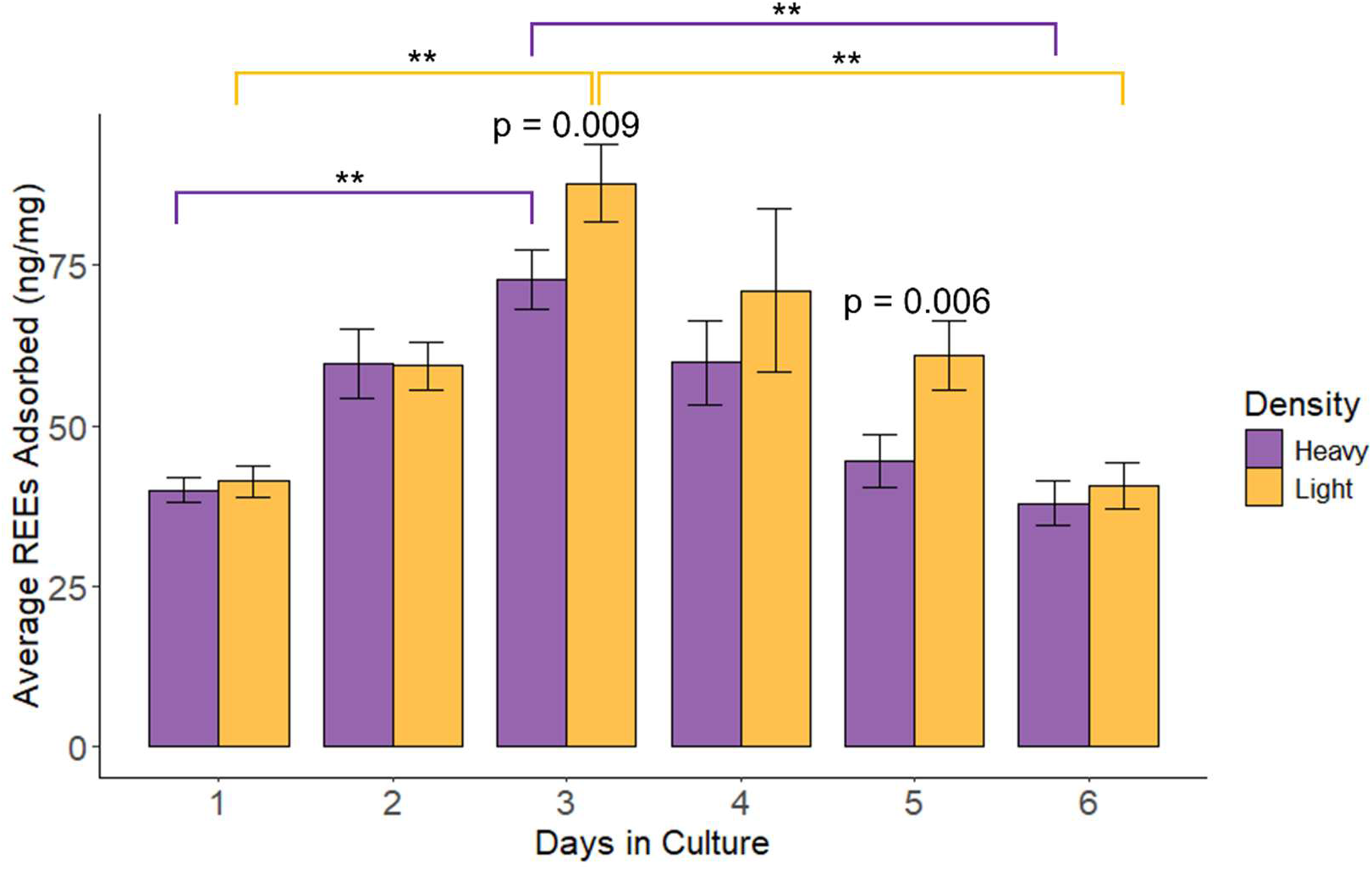
Average nanograms of light REEs and heavy REEs bound to cells normalized to dry pellet weight (mg) with statistical significance determined by paired student T-test showing significant differences between heavies and lights on days 3 and 5 and significant differences between days shown with brackets colored yellow for lights and purple for heavies.

Detailed analysis of individual light and heavy REEs shows consistent adsorption behavior among light REEs (Figure 2). The top five adsorbed light REEs were La, Ce, Nd, Sm, and Pr. Overall, the light REEs behaved similarly with the peak at Day 3 and then declining to Day 6. Neodymium had the highest adsorption with an average of 92 ng per mg of dry cells on Day 3. The top five adsorbed heavy REEs were Y, Gd, Tb, Dy and Ho. Gadolinium was the best adsorbed heavy REE with an average of 79 ng adsorbed per mg of cells on Day 3. The heaviest REEs, Yb and Lu, have a greater decrease in adsorption from Day 3 to Day 6 than the other heavy REEs. For all REEs, Day 4 had the highest variability (ranging from 34.6 ng of Lu per mg of cells to 92.7 ng of Nd per mg of cells) likely driven by the variability in pellet weight. One replicate from Day 4 weighed 50% more than the other two replicates. Overall, the best adsorption capacity for *S. oneidensis* was on Day 3 with an average total REEs of 1.18 mg/g of dried cells.

**Figure 2.**
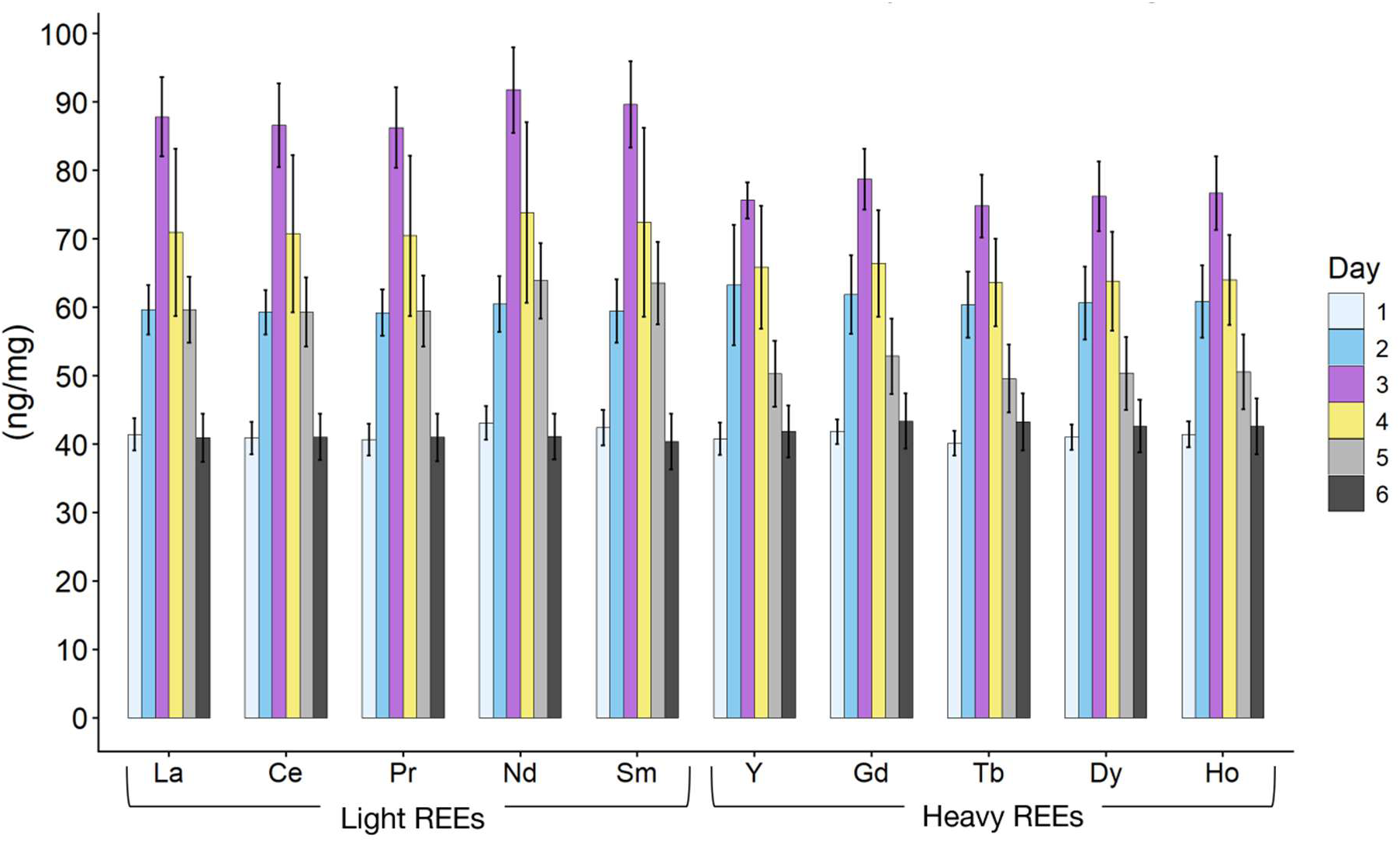
Top five light and top five heavy REEs (ng) adsorbed by *S. oneidensis* MR-1 normalized to dry cell pellet weight (mg)

These results are comparatively lower than other biosorbents. *Euglena mutabilis,* a single-celled protist, was found to have a lanthanum adsorption capacity of 7.7 mg/g.^29^ Freeze-dried *Galdieria sulphuraria* biomass pretreated with an acid solution achieved adsorption of 34.59 mg of yttrium per gram of biomass.^30^ Native *Arthrobacter nicotianae* was shown to have a Nd adsorption capacity of 69 mg/g, and *E. coli* genetically engineered to express lanthanide binding tags has an Nd adsorption capacity of 17 mg/g.^31^ A genetically engineered protein-based biosorbent made by fusing the lanmodulin from *Hansschlegelia quercus* to protein crystalline inclusions had an REE adsorption capacity ranging from 14 to 34.6 mg/g.^32^ The adsorption capacity of *S. oneidensis* may be improved by overexpressing the part of the organism responsible for binding REEs, which may be as simple as starving the bacteria to promote EPS formation.^33,34^ The observed difference between light and heavy REE biosorption (Figures 1 and 2) implies a specific biochemical or physiochemical mechanism rather than random accumulation. We, therefore, asked the question of where on the cell does REE binding occur, with the hypothesis that binding occurs in the EPS. To examine how the EPS contributes to REE adsorption, *S. oneidensis* was incubated with a standard REE solution, and the cells were separated from the loosely bound EPS (LB-EPS). The majority of both light and heavy REEs (∼80%) were bound to the LB-EPS regardless of the incubation time (Figure 3). There was no significant difference seen between light and heavy REE binding in the LB-EPS until Day 5 of the experiment. On Day 5 and continuing through Day 6, the LB-EPS bound significantly more heavy than light REEs.

**Figure 3.**
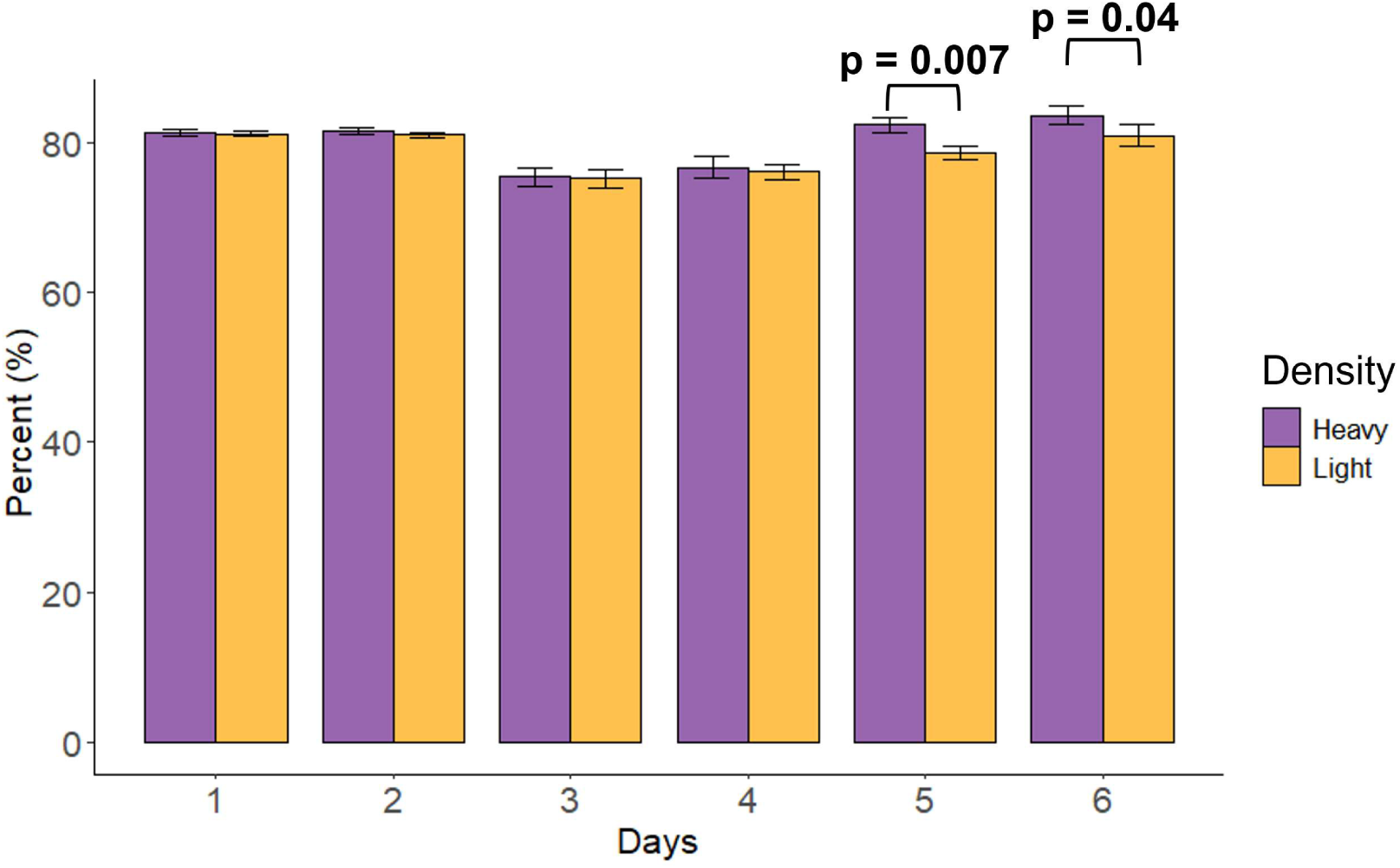
Percentage of LREEs and HREEs adsorbed to the loosely bound (LB) EPS after variable days in culture with REEs.

The EPS is a complex, multilayer matrix composed of polysaccharides, proteins, and DNA surrounding bacterial cells.^12,35^ EPS is rich in functional groups such as carboxyl, phosphoryl, and hydroxyl moieties that have strong affinity for multivalent cations, including REEs.^36^ Carboxyl and phosphoryl functional groups on cell surfaces have been shown to dominate REE biosorption to gram-negative bacteria.^37^ Surface complexation models have shown that LREEs best adsorb to phosphoryl group, and HREEs adsorb well to both carboxyl and phosphoryl groups.^38^ Furthermore, EPS from several organisms has been shown to bind a variety of metals. For example, EPS-covered *Pseudomonas aeruginosa* binds more Cr, Pb, and Cd than *P. aeruginosa* without EPS.^35^ Other studies have shown that REEs biosorb to *E. mutabilis* due to the abundant EPS in algal biofilms.^39^ Studies with *S. oneidensis* have shown that the EPS enhances extracellular electron transfer during the biotransformation of ferrihydrite, in part by binding Fe(II) and Fe(III).^12^ Our results show that the LB-EPS in particular is responsible for *S. oneidensis* binding of REEs.

The chemical composition of the EPS matrix of *S. oneidensis* is known to change in response to changing nutrient conditions.^40^ During the incubation with REEs, *S. oneidensis* was in stationary phase (Figure SI-2) in low nutrient media, and nutrient limitation is thought to result in more EPS production.^41^ The EPS is the primary component of biofilms. *S. oneidensis* biofilm development has been shown to be dependent on the *mxd* gene cluster, and expression of these genes is induced by carbon starvation.^42^ When grown in a nutrient dense broth like LB, *S. oneidensis* forms fragile clusters of cells weakly anchored and easily separated with minor agitation.^43^ Furthermore, the same study found that *S. oneidensis* formed stronger (i.e. more compact and cohesive) biofilms at 24 and 48 hours when given 0.5mM versus 2mM of lactate; however, after five days biofilms were identical regardless of the amount of lactate.^43^ Indicating that the EPS of *S. oneidensis* was of a similar composition in both scenarios by Day 5. That study also showed that *S. oneidensis* biofilms reached a quasisteady state after 5 days. The experiments conducted here provided one dose of 20mM Na-L-lactate which was not enough to stimulate growth but allowed for the maintenance of stationary phase (Figure SI-2). These results support the findings described above by showing that selectivity towards heavy over light REEs in the EPS became significant on Day 5, potentially because the cells entered a different state, a steady state, induced by starvation conditions.

While determining the chemical composition of the EPS is beyond the scope of this study, results imply that by Day 5, the composition of the LB-EPS had been modified to prefer adsorption of HREEs over LREEs (Figure 3). The LB-EPS of *S. oneidensis* typically contains more carboxyl groups and phosphoryl groups than the TB-EPS.^12^ The differences observed in the adsorption of HREEs and LREEs to *S. oneidensis* on Days 5 and 6 are likely due to changes in both the composition of the EPS and the physiochemical differences between light and heavy REEs that influence their binding affinity to EPS functional groups.

### 3.2 REE Desorption

After demonstrating the REE binding capacity of *S. oneidensis* and locating the majority of REEs on the LB-EPS, desorption of REEs from cells was tested using sodium citrate. Over 90% of total REEs could be desorbed using 0.5M sodium citrate (Figure 4), whereas desorption controls using only nanopure water desorbed only ∼5% of total REEs. After 2 hours, desorption of LREEs averaged 90% and HREEs averaged 85% (Figure 4). Within the first 10 minutes of incubation, 78% of LREEs and 72% of HREEs were desorbed. More LREEs were desorbed in the first 12 hours compared to the HREEs. However, after 12 hours, HREEs were more completely desorbed.

**Figure 4.**
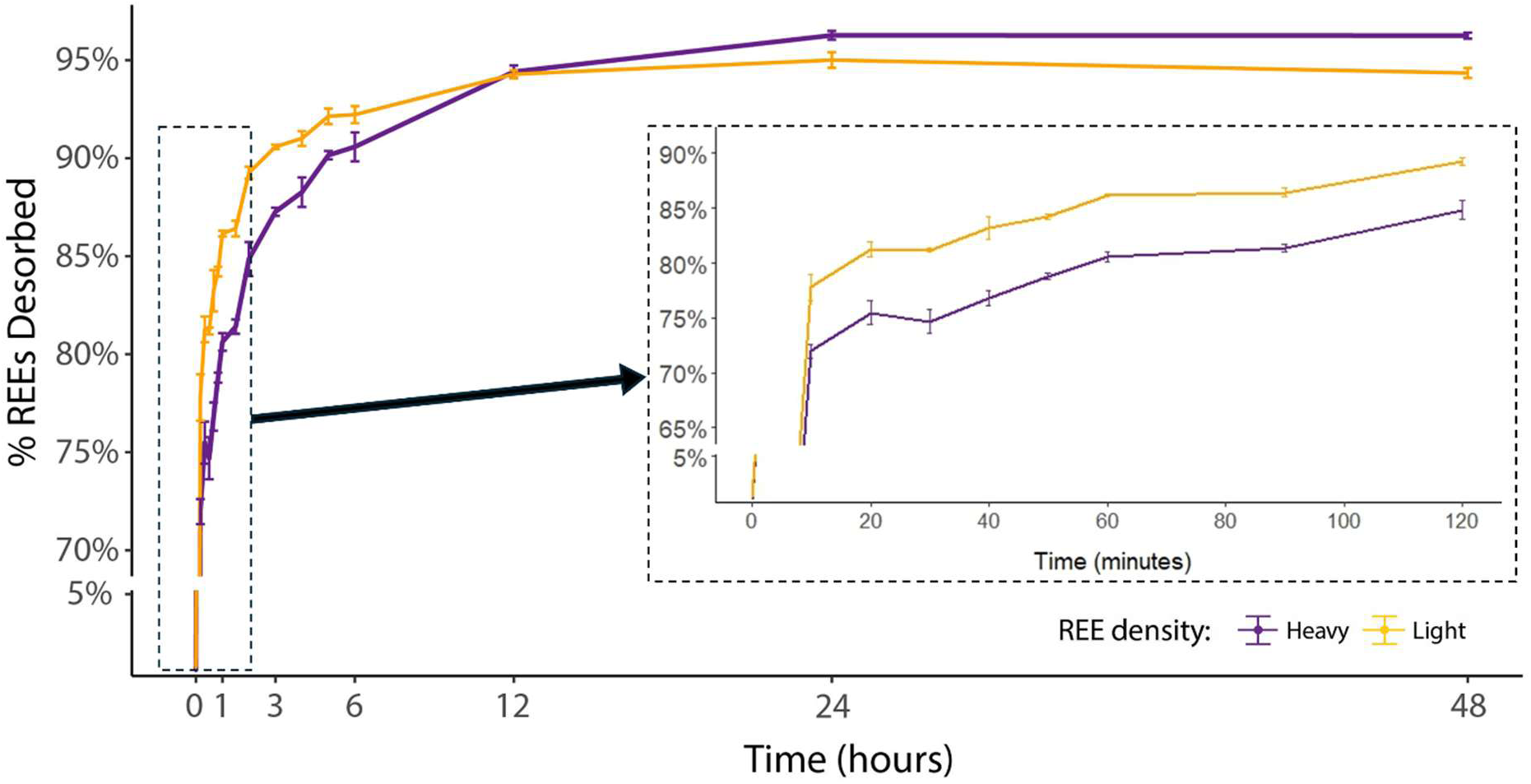
Percentage of heavy and light REEs desorbed over time from *S. oneidensis* with 0.5M sodium citrate. Inlay depicts HREE and LREEs desorbed in the first 2 hours of sodium citrate exposure.

Sodium citrate desorption results agree with other studies using different desorbants which also show desorption occurs faster than biosorption.^29^ For example, desorption of lanthanum bound to *E. mutabilis* for one hour using 0.1 M Na_2_EDTA at pH 5 had an equilibrium around 79%, with over 60% of that desorption occurring in the first 10 minutes.^29^ Our results showing that the difference in desorption between LREE and HREEs was never greater than 6% is in contrast to other studies using sodium citrate with different bacteria. For example, Yan and Chen (2024) showed that *E. coli* subjected to 0.05 M citrate at pH 6 desorbed HREEs more completely than LREEs after only 2 hours. This may be due to a difference in the location of REE binding on *E. coli* verses *S. oneidensis.* That same *E. coli* study showed that the EPS adsorbed only 31% of the REEs, but the cell wall was a major site of REE adsorption,^25^ whereas the work here showed ∼80% of REEs in the LB-EPS (Figure 3). Yan and Chen (2024) also found that *E. coli* adsorbed Yb, Lu, and Y better than any other REEs,^25^ whereas the work here with *S. oneidensis* showed better adsorption with light REEs (Figure 2).

We can infer that 20% of REEs are bound either in the TB-EPS or on the cell wall since ∼80% of REEs were located in the LB-EPS and preliminary work (Figure SI-3) indicated that REEs are not accumulating inside of the cells. Figure 4 shows that the first 10 minutes of desorption removes the majority of the REEs. The REEs released in the first 10 minutes are likely from the external layers of the EPS (primarily composed of LB-EPS), as those are the most accessible to sodium citrate and therefore acted on first. In our experiments, the 0.5M sodium citrate solution had a pH of 7.75. At that pH, the three carboxyl groups of each citrate anion are deprotonated and compete with functional groups of the EPS for REE binding to form REE-citrates. After 10 minutes, desorption slows and a different desorption mechanism may be occurring, or the remaining REEs are more strongly bound to the TB-EPS or cell surface than they were to the LB-EPS. One potential alternative mechanism is physical disintegration of the external layers of the EPS, which sodium citrate treatment has been shown to do in waste activated sludge.^44,45^ Studies on waste sludge bacterial communities have shown that at a pH of 6.7, citric acid has a destructive effect on external EPS and a relatively small impact on TB-EPS.^44,46^ The solubilization of proteins and polysaccharides in the LB-EPS may slowly expose more REEs that can then be competitively bound to citrate compounds. Sodium citrate treatment for longer periods of time (multiple days) or at an acidic pH (≤4) has been shown to disrupt all layers of the EPS and even cell membrane structure.^44^ We took OD measurements during a 2 day incubation of *S. oneidensis* cells in the 0.5M sodium citrate solution to test whether or not cell membranes were being destroyed over time.

Optical density measurements at 600 nm show that 2 days suspended in 0.5 M sodium citrate results in only a 1% reduction in cell density (Figure SI-4), albeit with a high degree of variability. Within this same time period, over 90% of REEs are desorbed from cells into solution. This is important because it shows that the majority of REEs can be recovered with less than 1% cell destruction, potentially permitting the reuse of cells for multiple rounds of REE adsorption and desorption. This also confirms that sodium citrate at circumneutral pH is causing minimal destruction to cell membranes. More work is needed, but an initial hypothesis is that cells may need time to regenerate EPS layers after desorption to restore REE adsorption capacity.

### 3.5 Desorption of Coal Extracted REEs

Building on previous findings that *S. oneidensis* MR-1 can extract REEs from coal,^14^ we evaluated the use of sodium citrate to desorb these elements from the bacterial biomass. To investigate this, *S. oneidensis* MR-1 was incubated with REE-containing coal at 10% w/v. After a week-long incubation, cells were separated from coal and culture media and then subjected to REE desorption using sodium citrate for 24 hours. Of the REEs extracted from coal and bound to the cells, the percent desorbed (68-86%) from those cells by sodium citrate is shown in Figure 5. Yttrium (Y), lanthanum (La), and cerium (Ce) were three of the most abundant REEs in the Alaskan coal. After one week in culture with 1.4g of coal, cells were able to adsorb 2.0 ng of Y, 1.5 ng of La, and 2.9 ng of Ce. Sodium citrate desorbed between 69-77% of the yttrium and over 80% of La and Ce from the surface of *S. oneidensis* (Figure 5).

**Figure 5.**
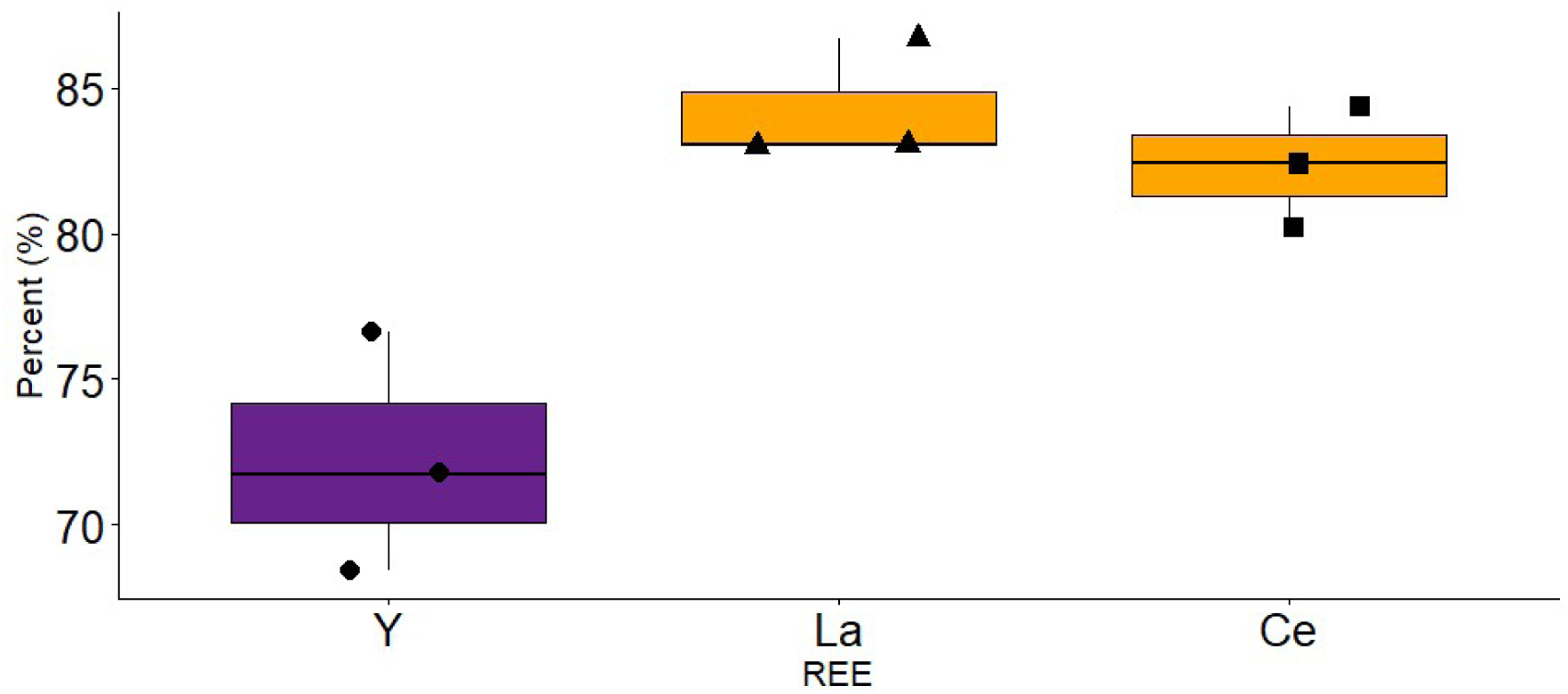
Three REEs extracted from Alaskan coal and desorbed from *S. oneidensis* in 24 hours with 0.5M sodium citrate.

Percent desorbed after 24 hours with the coal-extracted REEs (Figure 5) was lower than those desorbed after incubation with a standard solution of REEs (Figure 4). This may be explained by differences in the binding of REEs from a standard solution verses the binding of REEs from a mineral matrix. The REEs in this coal and others are known to be associated with the organic fraction.^47,48^ Organic matter compounds released from the coal during extraction may be involved in REE binding but are absent in REE binding from the defined standard REE solution experiments. The EPS is also known to have a role in the degradation of organic matter.^41^ By participating in the degradation of the coal or by the coal providing a surface for the bacteria to adhere to, the composition of the EPS may be different in the coal incubations verses the standard solution experiments. Overall, sodium citrate desorption is an effective recovery method for biomined REEs, but further research is needed into the extraction mechanism and its influence on REE binding to *S. oneidensis*.

## 4. Implications

The differences in light and heavy REE adsorption on Days 3 and 5 indicate the potential of *S. oneidensis* to be useful in biotechnological applications. Moreover, the results in Figure 3 strongly indicate that REEs are adsorbed to the LB-EPS of *S. oneidensis* cells which would allow for the reuse of cells for multiple rounds of adsorption and desorption. The observation that REEs are best adsorbed on Day 3 suggests a biological sweet spot where the physiological state of the cells, coupled with EPS maturity, leads to maximized REE adsorption. The differentiation between LREEs and HREEs in the EPS becoming significant on Day 5 indicates that as cells progress into late stationary phase or even death phase, the composition of the LB-EPS preferentially binds HREEs (Figure 4). *S. oneidensis* first binds almost all the REEs in solution with little discrimination, and then on Day 3 significantly more LREEs remain adsorbed than HREEs (Figure 1). This separation required no external stimulation or chemical additions. In particular, the faster desorption of heavy REEs (Figure 2) without the use of any desorbing agent could be useful for passive separation of REEs of different densities. Furthermore, the switch from more LREEs desorbed to more HREEs desorbed after 12 hours in sodium citrate solution indicates that LREEs may dominate the LB-EPS and HREEs may dominate the TB-EPS. This potential fractionation of LREEs and HREEs within layers of the EPS warrants a closer look at the biomolecules involved in REEs binding and further studies to identify the genetic engineering potential for the expression of those molecules. The selectivity of *S. oneidensis* bioadsorption and subsequent desorption could be improved by manipulating culture conditions to promote development of specific layers of the EPS.

This work complements the current literature on *S. oneidensis*’ capacity to both extract and adsorb REEs and improves the state of the science by describing a completely circumneutral pH bioprocess for REE extraction and concentration. Similar studies have depended on an acidic desorbant, such as Fujimoto et al., (2016) which demonstrated that *Shewanella putrefaciens* (a close relative to *S. oneidensis*) was able to extract REEs present in ferromanganese nodules and that upwards of 80% of an REE adsorbed to the cells, and then 70-80% of those REEs could be recovered by washing the cells three times with 0.01 M HCl.^16^ Moreover, Jiang et al. (2026) recently developed a bio-hybrid material composed of *S. oneidensis* MR-1 and microbially synthesized ferrous sulfide (FeS@MR-1) that synergistically adsorbed REEs from real mine wastewater with > 98 % efficiency and high selectivity over non-REE metals.^17^ Collectively, these works highlight *S. oneidensis* as a promising chassis for sustainable REE biomining, combining physiological adaptability, metal-reducing activity, and tunable surface chemistry for recovery and separation from diverse feedstocks including secondary sources and contaminated environments.

## Supplementary Information

**Figure SI-1:**
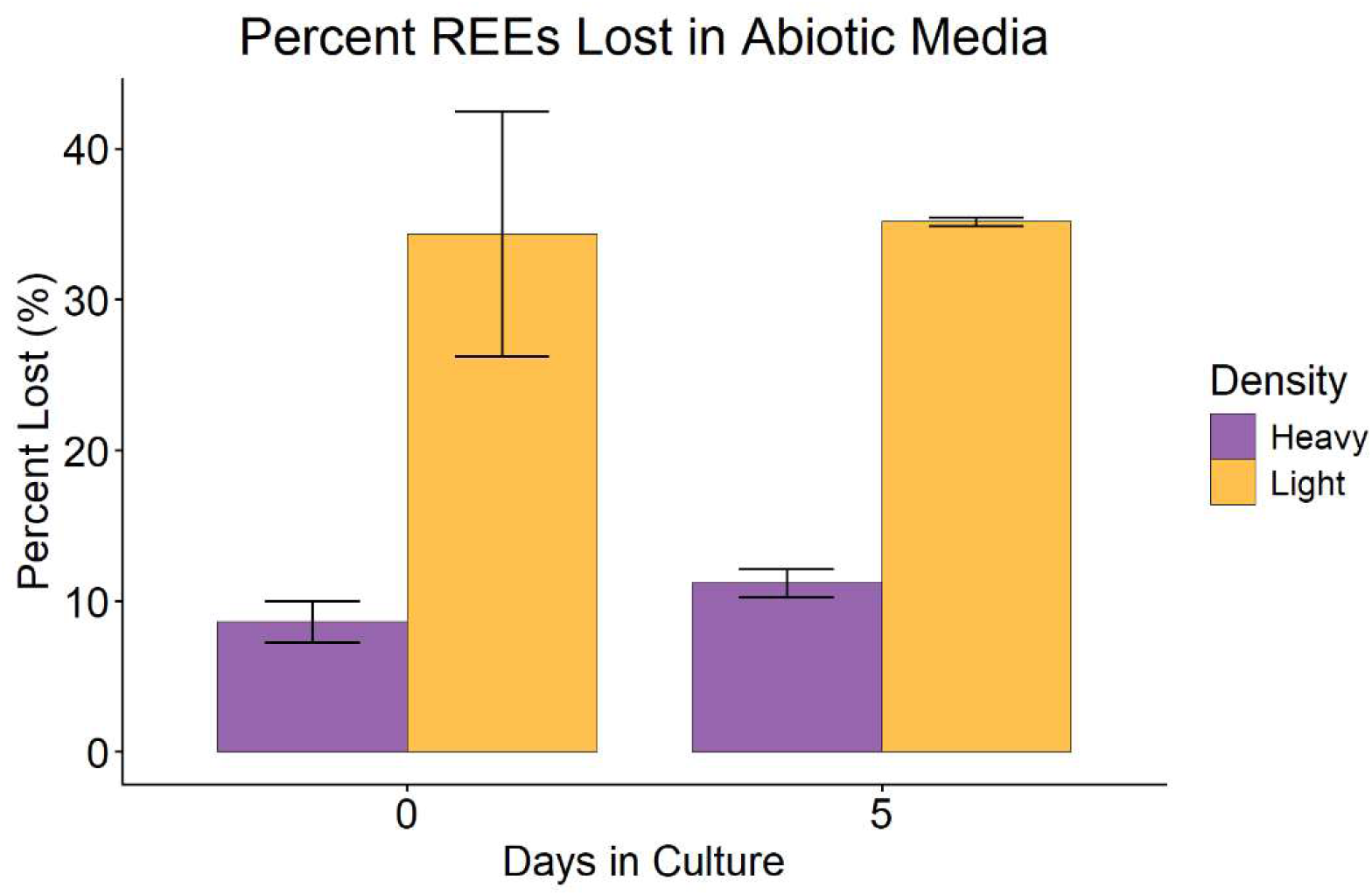
Average percent of light and heavy REEs lost from the media in abiotic conditions after zero and five days.

**Figure SI-2:**
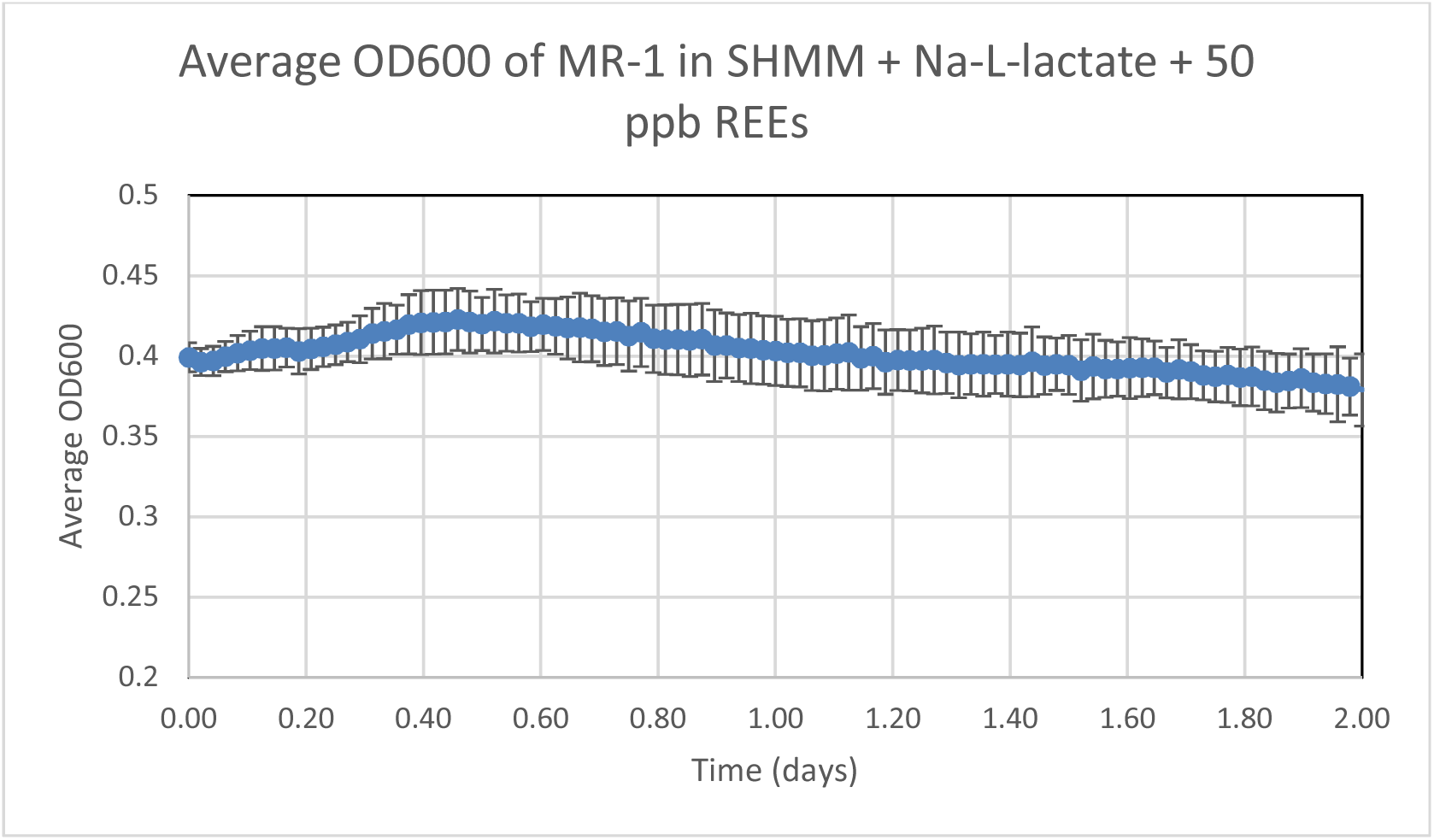
Growth curve for Shewanella oneidensis MR-1 in SHMM with 20mM Na-L-lactate and 50 ppb CMS-1 REE standard mixture over two days.

**Figure SI-3:**
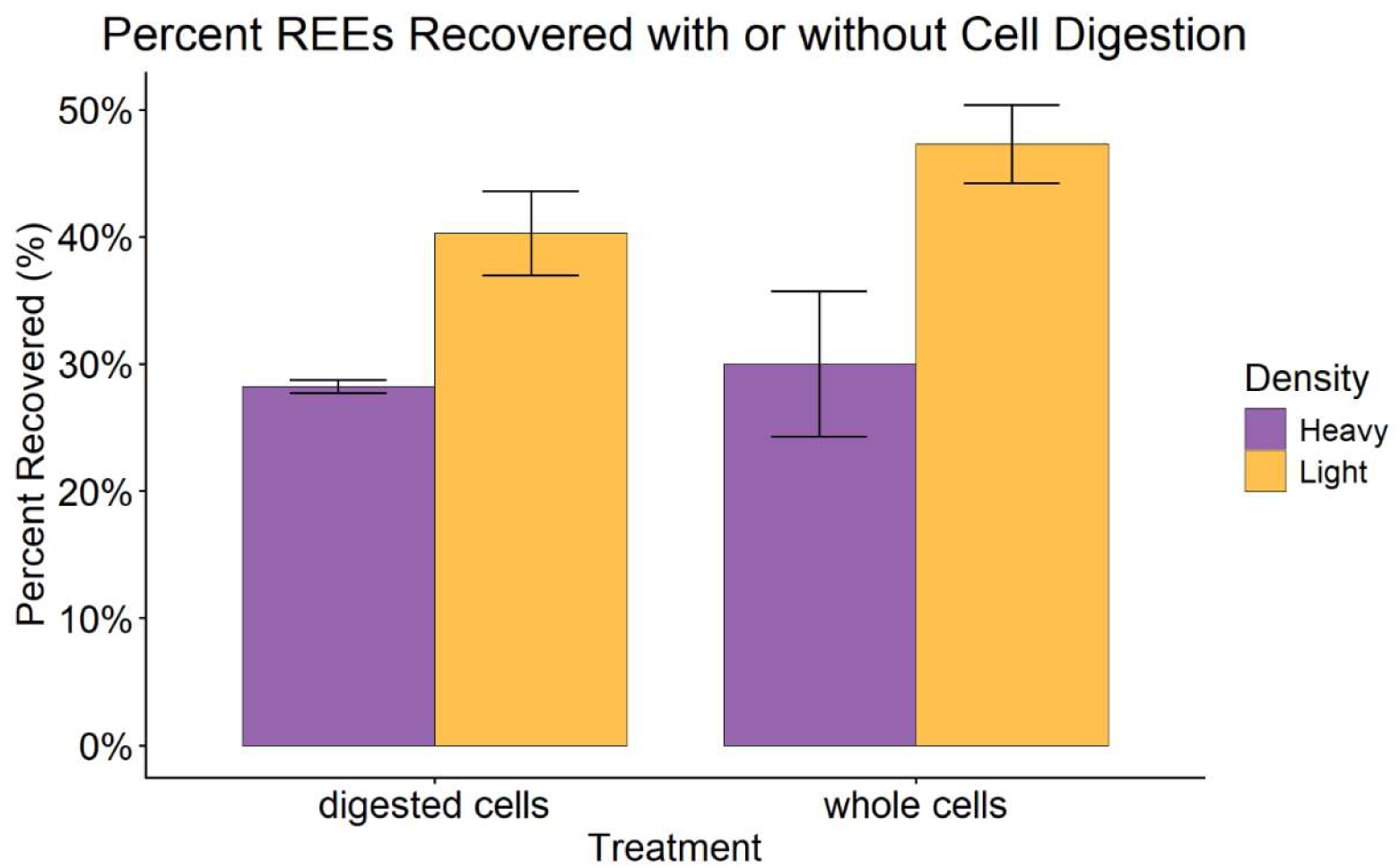
REEs recovered from a seven-day incubation of S. oneidensis with a standard solution of 50ppb of each REE. Cells were either acidified immediately after pelleting from culture (whole cells) or digested using proteinase K for 24 hours prior to acidification (digested cells).

**Figure SI-4:**
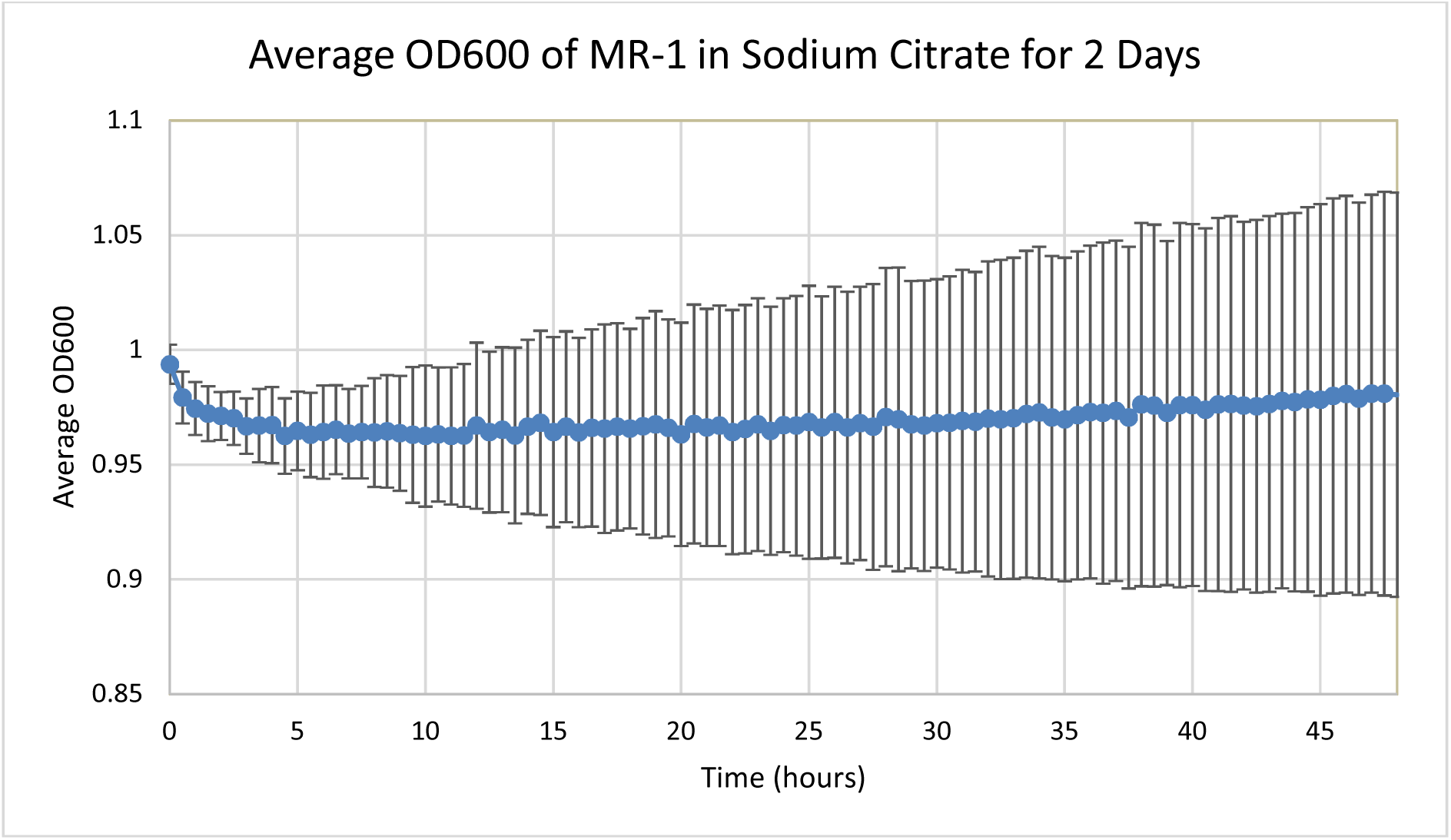
Growth curve for Shewanella oneidensis MR-1 concentrated from 3 ml of SHMM + L-lactate + CMS-1 to 1ml of 0.5M sodium citrate.

